# Title: Determinants of Minimum Acceptable Diet among 6–23 Months Age Children in Ethiopia: A Multilevel Analysis of The Ethiopian Demographic Health Survey

**DOI:** 10.1101/393678

**Authors:** Aberash abay, Dejen Yemane, Abate Bekele, Beyene Meressa

## Abstract

**Background:** Though infant and young children should be fed according to a minimum acceptable diet to ensure appropriate growth and development, only 7% of Ethiopian 6-23 months age children meet the minimum acceptable dietary standards, which is lower than the national target of 11% set for 2016. Therefore, this study aims to assess the individual and community level determinants of minimum acceptable diet among 6–23 months age children in Ethiopia.

**Methods:** This study analyzed retrospectively a cross-sectional data on a weighted sample of 2919 children aged 6-23 months nested within 617 clusters after extracting from Ethiopian Demographic and Health Survey 2016 via the link www.measuredhs.com. By employing bi-variate multilevel logistic regression model, variables which were significant at the p-value < 25 were included in multivariable multilevel logistic regression analysis. Finally, variables with p-value < 0.05 were considered as significant predictors of minimum acceptable diet.

**Results:** Only 6.1% of 6-23 months age children feed minimum acceptable diet in Ethiopia. Children 18-23 months age (AOR=3.7, 95%CI 1.9, 7.2), father’s with secondary or higher education (AOR=2.1, 95%CI 1.2, 3.6), Employed mothers (AOR=1.7, 95%CI 1.2, 2.5), mothers have access to drinking water (AOR=1.9, 95%CI 1.2, 2.9), mothers with media exposure (AOR=2.1 95%CI 1.1, 2.7) were positive individual level predictors. Urban mothers (AOR=4.8, 95%CI 1.7, 13.2)) and agrarian dominant region (AOR=5.6, 95%CI 2.2, 14.5) were community level factors that significantly associated with minimum acceptable diet of 6–23 months age children.

**Conclusion:** Both individual and community level factors were significantly associated with minimum acceptable diet of 6-23 months age children in Ethiopia, suggesting that nutritional interventions designed to improve child health should not only be implemented at the individual level but tailored to community context as well.

## Introduction

After 6 months, breast milk is no longer adequate to meet the nutritional needs and increasing demand of nutritional requirements of infants and children[1]. Complementary feeding is the process of transition from exclusive breastfeeding to other foods besides breast milk[2]. During this period timely introduction of complementary feeding with a variety of foods should be added to the child’s diet to ensure their nutritional requirement[3]. World Health Organization (WHO) has established guidelines for infant and young child feeding (IYCF) practices for 6–23 months age children by considering minimum acceptable diet (MAD) as one of the eight core indicators of complementary feeding [4]. It is the combination of minimum dietary diversity and minimum meal frequency. Minimum dietary diversity for breastfed and non breast feed children 6-23 months is defined as receiving four or more food groups out of the seven food groups[5] [7].

Minimum meal frequency for breastfed and non breast feed 6-23 month children is defined as two or more feedings of solid, semi-solid, or soft food for 6-8 months, three or more feedings for 9-23 months breast feed and four times for non breast feed children[5].

Low dietary diversity and meal frequency practices are determinant for health and growth in children less than 2 years of age. They increase the risk of under-nutrition, illness, and mortality in infants and young children [6]. Even with optimum breastfeeding, children will become stunted if they do not receive sufficient dietary diversity and frequency over 6 months of age [7]. Supplementing breast feeding with nutritious complementary foods can reduce stunting among children of this age by 20% [8].

According to the recent demographic and health survey reports of 10 Asian and African countries, including Ethiopia feeding a child minimum acceptable diet ranges from 7% in Ethiopia to 36% in Nepal. This indicates that, feed the minimum acceptable diet is a major problem both globally and in developing countries[1,3, 9-16].

The Ethiopian government has been implementing the national nutrition program, IYCF and has recently developed a multi-sectoral plan of nutrition intervention (Sekota Declaration), which aims to address the immediate, basic and underlying causes of malnutrition to end child under nutrition in Ethiopia by 2030 [17]. These interventions are meant to tackle the nutrition problems in children, including those with inappropriate feeding. However, the progress was not satisfactory, particularly; the MAD has increased from 3% to 7% in a decade (2005-2016). Factors associated with the minimum acceptable diet are complex, ranging from the individual to the community level factors. Though some studies conducted about determinants of complementary feeding in Ethiopia, no information was documented about minimum acceptable diet independently. However, these small scale studies were limited in scope as the data used were not nationally representative. In addition, they are also limited in their methodology in which they have applied traditional logistic regression to identify aforementioned predictors of minimum acceptable diet, using a traditional level logistic regression analysis to analyze a data that has a hierarchical structure (i.e. Children are nested within the communities) violates the independence assumptions of regression and gives an incomplete picture to understand the true association of minimum acceptable diet and its determinants. Likewise, findings from such studies could not be generalized to the entire Ethiopian children.

Hence, to address these limitations and to further document the significant effect of individual and community level factors on minimum acceptable diet, this study utilized a multilevel logistic regression model. Therefore, the purpose of this study will be to determine the individual and community level factors that are associated with minimum acceptable diet in Ethiopia.

## Methods and materials

### Data source and study subjects

For this study, the EDHS 2016 child recode data were accessed from www.measuredhs.com. It is the recent and the fourth nationally representative survey data from 9 regions and two administrative cities. The survey used a two-stage cluster sampling design with rural-urban and regions as strata yielding 21 sampling strata. In the first stage, a total of 645 enumeration areas (EAs) was selected. An EA is a geographic area consisting of 200-300 household, which served as a counting unit for the census. In the second stage, a fixed number of 28 households per cluster were selected randomly from the household listing. A total of 16,650 eligible households and 15,683 women (15-49 years) were interviewed, making up response rates of 98% and 95%, respectively. All women who had at least one child living with them who was last born two-years preceding the survey were asked questions about the types of food the child had consumed during the day or night before the interview. Mothers who had more than one child within the two years preceding the survey were asked questions about the youngest child living with them [1].

In our study, a total of 2919 children of 6-23 months age who had measurement on their feeding behavior and who live with their mother during the survey were included. Accordingly, children who did not live with their mother during the survey (n=26), and those children’s who had incomplete information about MAD (n=98) were excluded from our study. The child recode data used in this study comprised all the data related with children and their parents. Further, the details of sampling design, data collection, and data quality are available in the EDHS 2016 reports [1].

### Study variables

The DHS recode-6 manual and questionnaire at the end the EDHS 2016 report along with other relevant literatures were used to select appropriate variables for current analysis. The outcome variable was minimum acceptable diet (MAD). It is a binary outcome variable and coded as 0 if the child didn’t feed minimum acceptable diet and coded as 1 if the child feed a minimum acceptable diet. The child is said to be fed with MAD if he/she had both minimum meal frequency and dietary diversity in both BF and non-BF children (39). The children who received solid, semi-solid or soft foods, two times for breastfed infants 6–8 months, three times for breastfed children 9–23 months and four times for non-breastfed children is said to have minimum meal frequency. The children with 6–23 months of age who received foods from four or more food groups of the seven food groups (Cereals, Legumes and nuts, Dairy products, Eggs, Flesh foods, Vitamin A rich fruits and dark green leafy vegetables and other fruits) are said to have minimum dietary diversity [2].

In this study, the effect of two-level explanatory factors i.e. the individual and community level factors was examined. The relationship between the explanatory variables and the minimum acceptable diet is depicted using the socio-ecologic model [18]. According to this model, individual dietary or health behavior is not only determined by the individual’s characteristics, but also the community level attributes (**Fig 1**). The individual level variables are further categorized as paternal and household characteristics, and child related variables. While, at community level, place of residence (either urban vs. rural or by agro-ecologic regions) and community level aggregate variables such as community poverty, women’s education, media exposure and ANC utilization were included. The EDHS did not capture data that can directly describe the characteristics of the community/clusters. Hence, we created community variables by aggregating the individual mothers’ characteristics within their clusters. The aggregates were computed using the average values of the proportions of women in each category of a given variable. Likewise, based on national median values the aggregate values were categorized into groups.

### Statistical analysis

The data were analyzed using STATA version 12. A sampling weight was used for computing all descriptive statistics to adjust for the non-proportional allocation of the sample to different regions and their urban and rural areas and for the possible differences in response rates. Hence, the actual representativeness of the survey results at both the national and regional levels is ensured. The ‘Svy’ command was used to allow for adjustments for the cluster sampling design. Tables were used for data presentation. Frequency and percentage were used to report categorical variables.

To examine the effect of multilevel factors on the individual dietary behavior, multilevel modeling approach was used. The nested nature of EDHS data makes the use of traditional regression methods inappropriate because of the assumption of independence among individuals within the same group, assumption of equal variance across groups which are inherent in traditional regression methods are violated. Therefore, in this study, a two-level mixed effect logistic regression analysis was employed in order to estimate both independent (fixed) effects of the explanatory variables and community-level random effects on feeding MAD. The first level represents the individual (children) and the second level is the cluster (community). Hence, the log of the probability of feeding MAD was modeled using two-level multilevel model as follows:

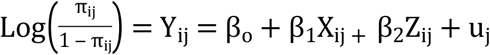

Where, *y_ij_* Is the feeding of MAD for the individual I in j cluster/community; X_ij_ and Z_ij_ are individual and community level variables for the i^th^ individual in group j, respectively. The β’s are fixed coefficients indicating a unit increase in X can cause a β unit increase in probability feeding MAD. While, the β_o_ is intercept that is the effect on feeding MAD when he effect of all explanatory variables absent (38). The uj shows random effect (effect of the community on mother’s decision to provide MAD) for the j^th^ community. By assuming each community has different intercept (β_0_) and fixed coefficient (β), the clustered data nature and the within and between community variations were taken in to account. The measures of community variation (random-effects) was estimated as intra class correlation coefficient (ICC) which is proportion of the total variance in feeding MAD due to variables operating at a community level. Hence, the ICC was estimated as: ρ = (σ^2^_*u*_ / (σ^2^_*u*_ + π^2^ /3)), Where, ρ is the ICC, σ^2^_*u*_ is the variance at the community level; π^2^ /3 = 3.29 represent the fixed individual level variance (37). Given the above assumptions, four models were developed, namely: Null-model – is a model with no explanatory variables; Model I – is a model with only individual level factors are controlled; Model II – a model with only community level factors are controlled and Model III – is also called combined model which controls the effect of both individual and community level explanatory variables on mothers decision to provide MAD to their child (**Table 1**).

The null model – the model with no covariates – was fitted to examine the between community variation and to justify the use multilevel analysis. Likewise, about 40% of the total variance (ICC) in the odds of feeding a child with MAD is due to community level factors (P<0.001), indicating using multilevel modeling is better to get valid estimates than using ordinary logistic regression. The Proportional Change in Variance (PCV) was computed for each model with respect to the empty model to understand the relative contributions of both individual and community level variables to the community level variance on feeding MAD to a child. It was calculated as PCV = (Ve-Vmi)/Ve; where, Ve is variance in the empty model and Vmi is variance in successive models (**Table 1**).

The between-community variability declined in successive models, from 39.6% in null model, to 28.3% in individual-level only model, 27.5% in community-level only model and 26.6% in combined model. Akaike information criterion (AIC) was used to select a best model that explains the variation in feeding MAD well. The model with the lowest AIC is the preferred, accordingly the combined model showed lowest AIC than others. The combined model implied that 44.9% of the total explained variance in the odds of feeding MAD to a child could be attributed to both individual and community level characteristics in the model. However, the variance in the combined model remained significant (p<0.001), indicating the presence of other community level factors – which were not included in our model – that can explain the feeding of MAD to a child. In other words, even though the unexplained community level variance was reduced in the mixed model, the remaining community-level variance still remains significant (**Table 1**).

**Table 1:** Individual and community-level variances for multilevel random intercept Logit models predicting feeding 6-23 months age children with minimum acceptable diet in Ethiopia 2016.

| Random effect | Null model | Model I | Model II | Model III (combined) |
| --- | --- | --- | --- | --- |
| Variance | 2.16* | 1.30* | 1.25* | 1.19* |
| ICC (%) | 39.6 | 28.3 | 27.5 | 26.6 |
| PCV (%) | Reference | 39.8 | 42.1 | 44.9 |
| AIC | 1296.4 | 1153.3 | 1238.7 | 1142.7 |

Note: Null model – a model with no covariates; Model I – only individual level explanatory variables were included; Model II – only community level explanatory variables were included; Model III (Combined) – both individual and community level variables were included; \*significant at $p < 0.001$ ; AIC: Akaike Information Criterion.

The measures of association (fixed-effects) between the odds of feeding a child with MAD and various explanatory variables were expressed as adjusted odds ratio (AOR) at their 95 % confidence intervals (CI). The statistical significance was set at p-value of 0.05. The multicollinearity among independent variables was checked using variance inflation factor (VIF). The VIF value of all predictor variables was less than 10 indicating the absence of a significant correlation among the explanatory variables. The presence of interaction among the explanatory variables was checked and there was no significant interaction between them.

This study was approved by the Ethical Review Committee of College of Health Sciences, Mekelle University . Then, a written inquiry to access data from the measure DHS website was submitted to the ORC macro INC, Chicago and permission was obtained accordingly. This study was based on the EDHS 2016 dataset with all participants’ identifiers removed, no further effort was made to trace back the subjects and the data was kept confidential as per the agreement made with the ORC macro. Otherwise, the DHS was approved by the Ethiopian Health Nutrition and Research Institute (EHNRI) Review Board and the National Research Ethics Review Committee (NRERC) at the Ministry of Science and Technology, Ethiopia.

## Results

### Individual level characteristics of the study participants

Five hundred fifty-six (19%) children were in the age group 6–8 months, 492 (17%) in age 9-11 months, 1071 (37%) in 12–17 months and the rest 799 (27%) found in 18–23 months. More than half of the children (52.9%) were females. More than two-third 1942 (67%) children were born within more than 24 months of pregnancy interval. About three-fourth mothers 2110 (72%) were in age between 20-34 years. Sixty-two percent of the mothers and 1207 (44%) of fathers had no education. About forty percent of participants were Muslims followed by Orthodox 998 (34.2%). Regarding the employment status, 1685 (58 %) of the mothers were not working at the time of EDHS 2016 survey. More than three-fourth 2521 (86.4%) of households were headed by males. More than two-third 2009 (69%) of the households have at least five family sizes and 1387 (47.5%) were with two under five children. More than half of households 1519 (53%), travel 30 minutes or longer round trip to fetch drinking water. On the other hand, only 90 (3%) were used efficient cooking fuel (electricity) but most 2373 (81.9%) of households use wood as cooking fuel. The proportions of mothers with no ANC visit and at least 4 visits during their last pregnancy were nearly the same, 1019 (35%) and 1002 (34.5%), respectively (**Table 2**).

**Table 2:**
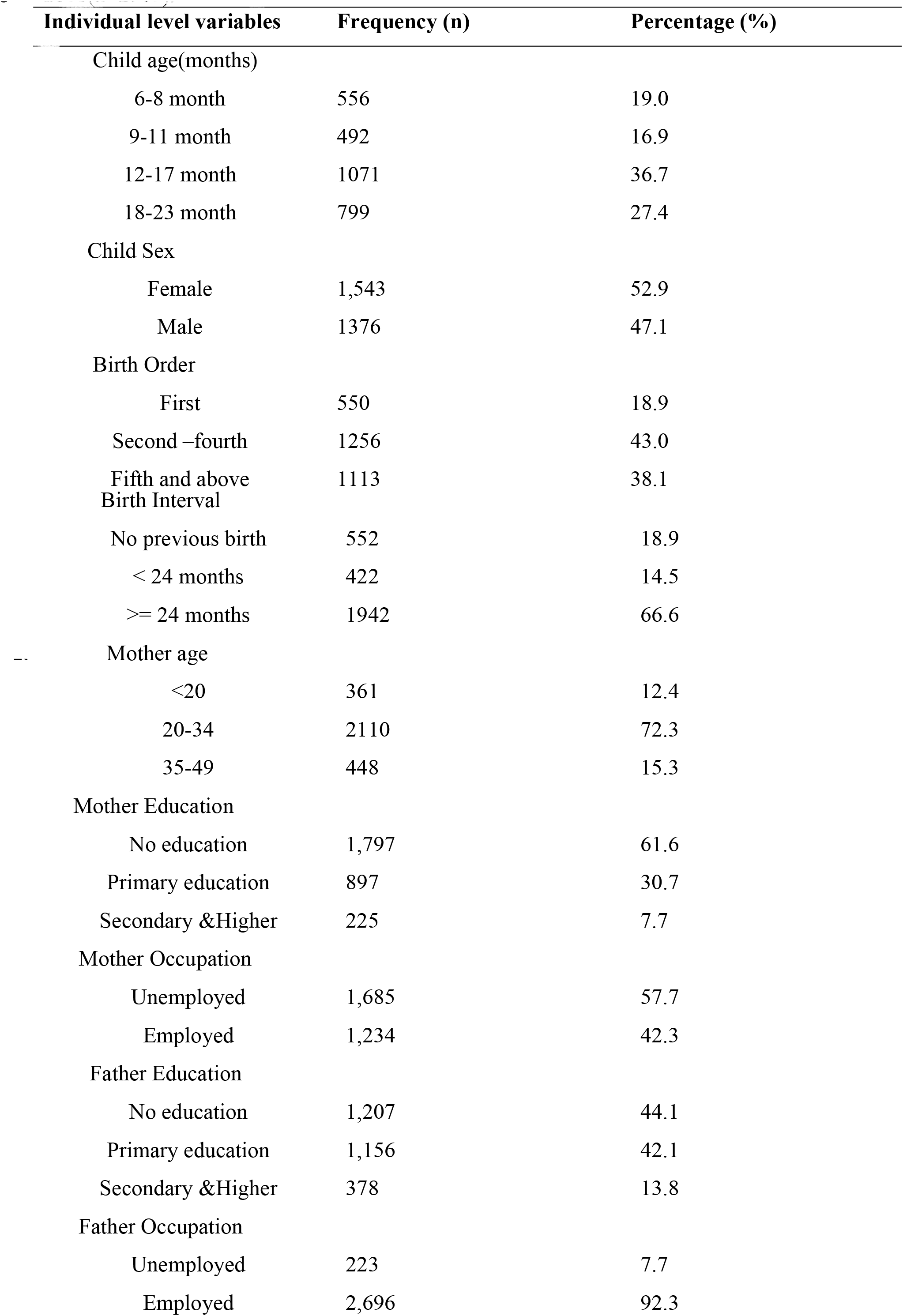
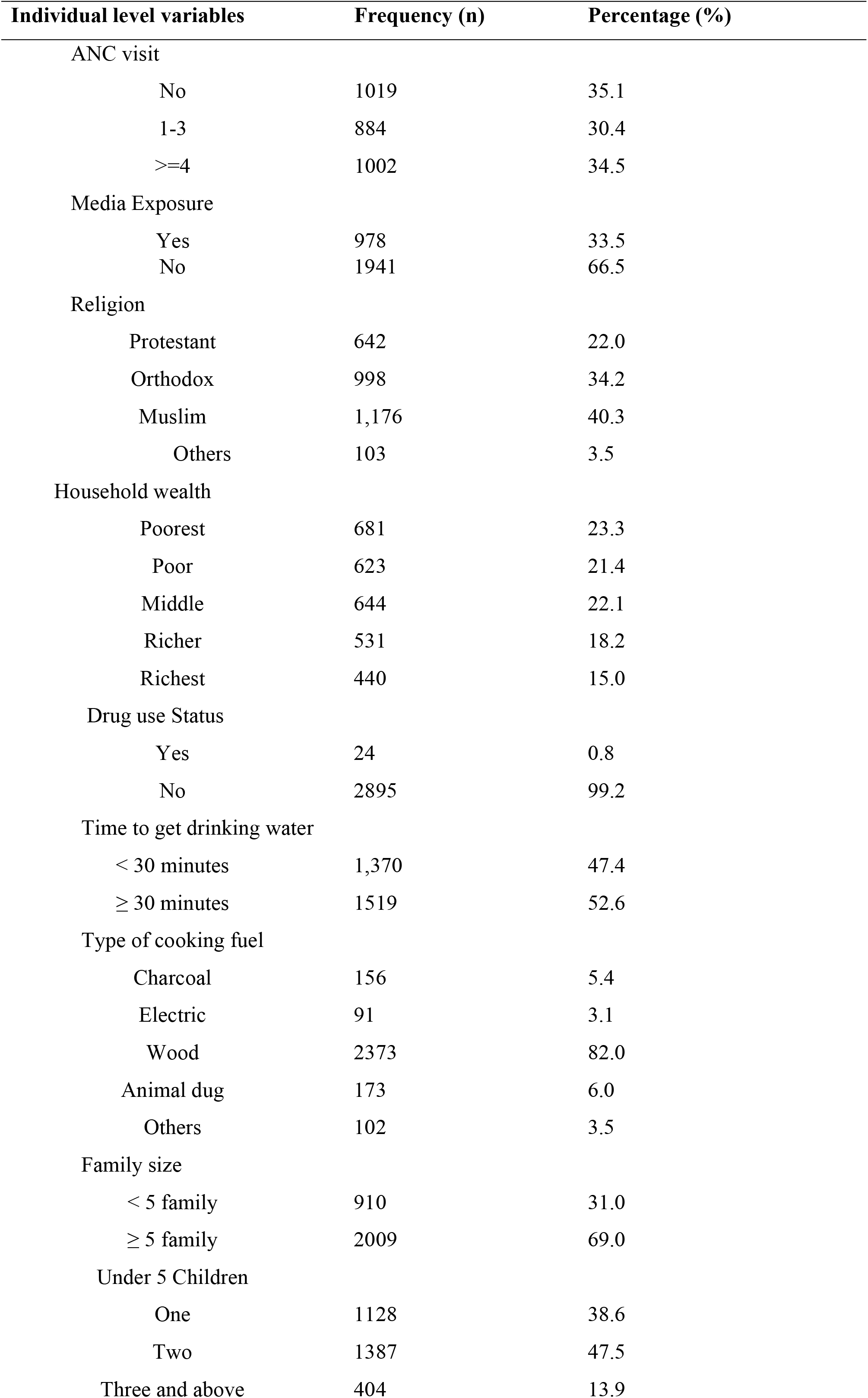
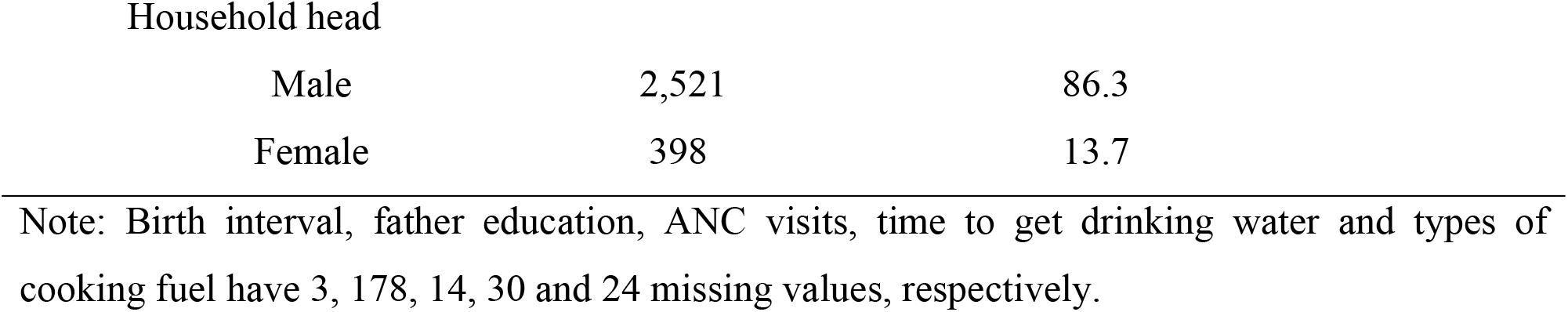
The individual level characteristics of 6-23 months age children in Ethiopia, EDHS 2016(n=2919).Note: Birth interval, father education, ANC visits, time to get drinking water and types of cooking fuel have 3, 178, 14, 30 and 24 missing values, respectively.

| Individual level variables | Frequency (n) | Percentage (%) |
| --- | --- | --- |
| Child age(months) |  |  |
| 6-8 month | 556 | 19.0 |
| 9-11 month | 492 | 16.9 |
| 12-17 month | 1071 | 36.7 |
| 18-23 month | 799 | 27.4 |
| Child Sex |  |  |
| Female | 1,543 | 52.9 |
| Male | 1376 | 47.1 |
| Birth Order |  |  |
| First | 550 | 18.9 |
| Second –fourth | 1256 | 43.0 |
| Fifth and above | 1113 | 38.1 |
| Birth Interval |  |  |
| No previous birth | 552 | 18.9 |
| < 24 months | 422 | 14.5 |
| >= 24 months | 1942 | 66.6 |
| Mother age |  |  |
| <20 | 361 | 12.4 |
| 20-34 | 2110 | 72.3 |
| 35-49 | 448 | 15.3 |
| Mother Education |  |  |
| No education | 1,797 | 61.6 |
| Primary education | 897 | 30.7 |
| Secondary & Higher | 225 | 7.7 |
| Mother Occupation |  |  |
| Unemployed | 1,685 | 57.7 |
| Employed | 1,234 | 42.3 |
| Father Education |  |  |
| No education | 1,207 | 44.1 |
| Primary education | 1,156 | 42.1 |
| Secondary & Higher | 378 | 13.8 |
| Father Occupation |  |  |
| Unemployed | 223 | 7.7 |
| Employed | 2,696 | 92.3 |

**Table 2:** continued
| Individual level variables | Frequency (n) | Percentage (%) |
| --- | --- | --- |
| ANC visit |  |  |
| No | 1019 | 35.1 |
| 1-3 | 884 | 30.4 |
| >=4 | 1002 | 34.5 |
| Media Exposure |  |  |
| Yes | 978 | 33.5 |
| No | 1941 | 66.5 |
| Religion |  |  |
| Protestant | 642 | 22.0 |
| Orthodox | 998 | 34.2 |
| Muslim | 1,176 | 40.3 |
| Others | 103 | 3.5 |
| Household wealth |  |  |
| Poorest | 681 | 23.3 |
| Poor | 623 | 21.4 |
| Middle | 644 | 22.1 |
| Richer | 531 | 18.2 |
| Richest | 440 | 15.0 |
| Drug use Status |  |  |
| Yes | 24 | 0.8 |
| No | 2895 | 99.2 |
| Time to get drinking water |  |  |
| < 30 minutes | 1,370 | 47.4 |
| ≥ 30 minutes | 1519 | 52.6 |
| Type of cooking fuel |  |  |
| Charcoal | 156 | 5.4 |
| Electric | 91 | 3.1 |
| Wood | 2373 | 82.0 |
| Animal dug | 173 | 6.0 |
| Others | 102 | 3.5 |
| Family size |  |  |
| < 5 family | 910 | 31.0 |
| ≥ 5 family | 2009 | 69.0 |
| Under 5 Children |  |  |
| One | 1128 | 38.6 |
| Two | 1387 | 47.5 |
| Three and above | 404 | 13.9 |
| Household head |  |  |
| Male | 2,521 | 86.3 |
| Female | 398 | 13.7 |

### Community level characteristics of the study subjects

More than nine in ten mothers 2679 (91.8%) were from the Agrarian dominant region. More than half 1937 (66.4%) of the mothers were from the community with high poverty. More than half mothers 1506 (52%) were from the low media access community. Most of the mothers 2576 (88.3%) were from rural areas and 2008 (69%) of the mothers were from the community with high ANC utilization (**Table 3**).

**Table 3:** The Community level characteristics of 6-23 months age children in Ethiopian, EDHS 2016 (n=2919)

| Community level | Frequency (n) | Percentage (%) |
| --- | --- | --- |
| Residence |  |  |
| Urban | 343 | 11.7 |
| Rural | 2576 | 88.3 |
| Region |  |  |
| Pastoralist dominant | 144 | 4.9 |
| Agrarian dominant | 2679 | 91.8 |
| City dwellers | 96 | 3.3 |
| Community poverty |  |  |
| High | 1937 | 66.4 |
| Low | 982 | 33.6 |
| Community women education |  |  |
| Low | 2053 | 70.3 |
| High | 866 | 29.7 |
| Community media exposure |  |  |
| Low | 1506 | 51.6 |
| High | 1413 | 48.4 |
| Community ANC utilization |  |  |
| Low | 911 | 31.0 |
| High | 2008 | 69.0 |

### Minimum acceptable diet feeding practice

In this study, of the total of 2919 children 6-23 months of age, only 6.1% were fed MAD. On the other hand, the proportion of children with minimum dietary diversity and minimum meal frequency were 11% and 42%, respectively (**Table 4**).

**Table 4:** Minimum meal frequency, Dietary diversity and minimum acceptable practice among children 6-13 months of age in Ethiopia, 2016 (n=2919).

| Diets | Frequency (n) | Percentage (%) |
| --- | --- | --- |
| Minimum acceptable diet |  |  |
| No | 2742 | 93.9 |
| Yes | 177 | 6.1 |
| Meal frequency |  |  |
| No | 1697 | 58.1 |
| Yes | 1222 | 41.9 |
| Dietary diversity |  |  |
| No | 2588 | 88.7 |
| Yes | 331 | 11.3 |

### Individual and community level determinants of feeding MAD

The individual and community level determinants of feeding MAD are presented in Table 5. At individual level: child age, father’s education, mother’s occupation, time to get drinking water, media exposure; and at community level: region of residence were independent predictors of feeding the child MAD.

The odds of feed the child MAD was nearly 4 times (AOR=3.7; 95%CI 1.9, 7.3) higher among children in age between 18-23 months than children at age 6-8 months. The odds of feeding the child with a minimum acceptable diet were 70% (AOR=1.7; 95%CI 1.2, 2.5) higher among employed mothers compared with unemployed mothers.

Mothers who have media exposure had nearly 2 times (AOR=1.95; 95%CI 1.3, 2.9) higher odds of feeding a MAD for their children than mothers with no media access. The odds of feeding MAD to a child was twice (AOR=2.0; 95%CI 1.3, 3.1) higher among mothers who travel less than 30 minutes to fetch drinking water. Father’s education was significantly positively associated with feeding minimum acceptable diet to the child. Fathers who educated secondary or higher had 2.4 times (AOR=2.4; 95%CI 1.4, 4.0) higher odds to feed their children MAD.

The region was a significant predictor of feeding the child MAD. Mothers from agrarian region had 5 times (AOR=5.1; 95% CI 2.0, 13.1) and city dwellers had nearly 5 times (AOR=5.4; 95%CI 1.9, 14.8) higher odds to feed their children MAD respectively compared with mothers from pastoralist (**Table 5**).

**Table 5:**
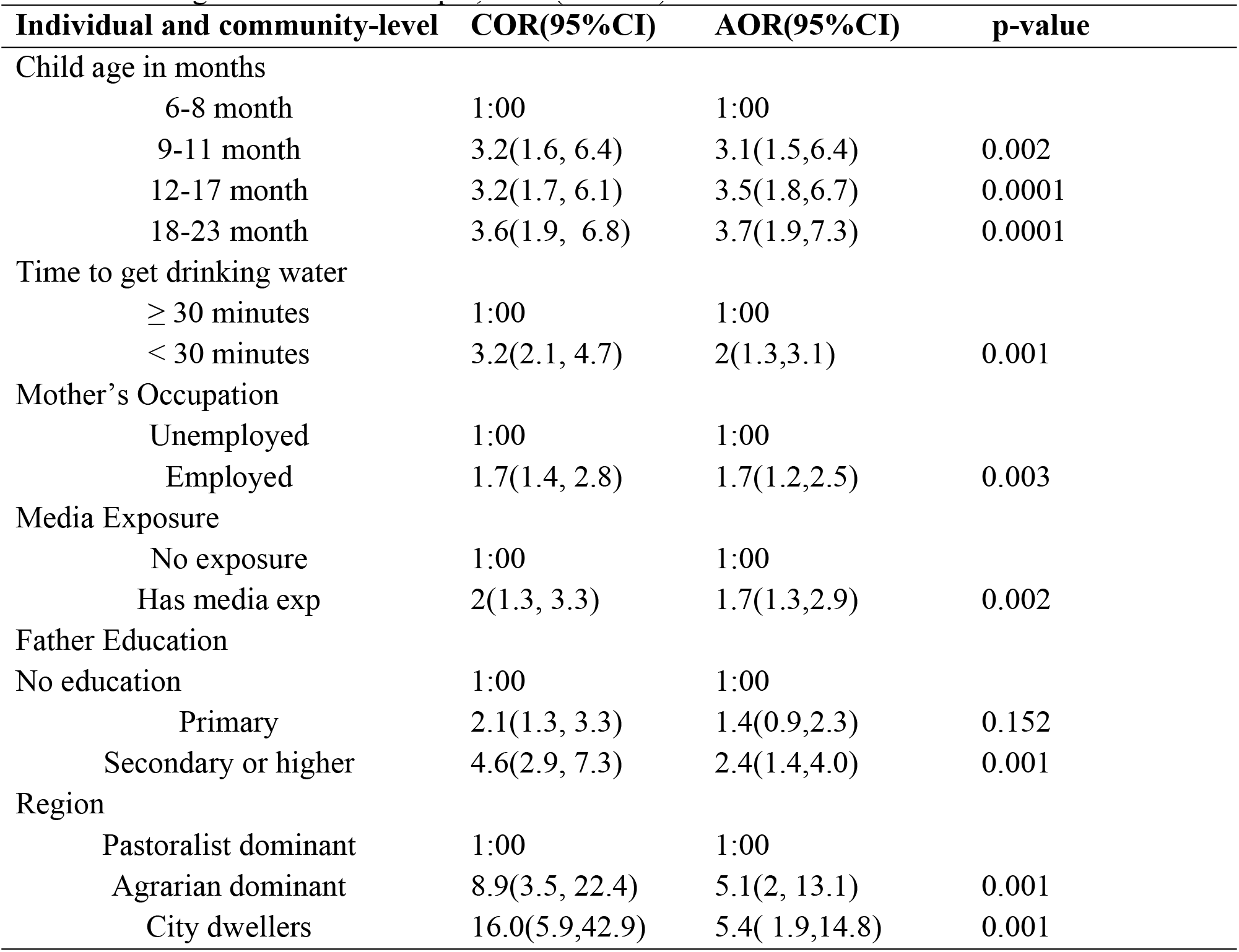
Individual and community level determinants of feeding minimum acceptable diet for 6-23 months age children in Ethiopia, 2016(n=2919).

## Discussion

This study used multilevel logistic regression analysis to address these two level factors. Accordingly the individual level factors such as the child’s age, mother’s occupation, father’s education, access to drinking water and media exposure showed a significant association with feeding minimum acceptable diet.

The age of the child was positively associated with feeding minimum acceptable diet for 6-23 months age children. Children with age 18-23 months were found to be significantly higher odds to be feed minimum acceptable diet than 6-8 months. This finding is consistent with evidences from Indonesia, Pakistan, Ghana and Uganda [7, 8, 19, 20]. This might be due to late introduction of complementary feeding and when they start complimentary feeding on time; they included only milk or cereal products. Other possibility could be mothers may perceive that younger the children, the poor ability of intestine to digest certain foods like banana, egg, pumpkin, carrot, green vegetables and Meat [21]. This could be further justified by traditional beliefs and practices, during introducing complementary food to infants in rural community, infants may develop diarrhea due to poor hygienic condition, but mothers could associate this problem with taking different food items and eventually she might not permit the child to taste unfamiliar foods. It can also be attributed to the feeding interest of the child too.

Father’s education level was significantly and positively associated with feeding the child minimum acceptable diet. Other studies in South Asia, Bangladesh and Nepal had reported a similar finding indicating that children whose fathers had secondary or higher level of education were more likely to be provided with the recommended acceptable diet as compared to the children whose fathers did not have any education [22, 23, 24]. This could be due to educated fathers were more likely to have information (media exposure), understand educational messages about child feeding easily, might have received lessons on child feeding in the curricula at school that would increase their knowledge about the importance of child feeding. However, different finding was reported from study done in Serilanka, in which paternal education was not associated with any of complementary feeding indicators [25]. This difference could be due to study area and sample size. The current study included large population from different geographic regions with various culture, beliefs, and traditions.

Mother’s occupational status was also significant predictor of feeding minimum acceptable diet to the child. Children who have employed mothers were more likely to be feed minimum acceptable diet. This result is consistent with study conducted in Indonesia and Serilanka [19, 25]. It could give us an indication that mother’s earning ability is an important factor in feeding the child a minimum acceptable diet in Ethiopia. Enhanced access to resources, wider social networks and growing understanding of their social environment could increases opportunities to feed the child minimum acceptable diet than unemployed mothers.

This study also revealed that time to get drinking water showed a positive effect on feeding the children a minimum acceptable diet. Mothers who travel less than 30 minutes round trip to fetch drinking water had higher odds to feed their children a minimum acceptable diet than those who travel 30 or more minutes. This could be attributed to that, mothers who travel less than 30 minutes to fetch drinking water could have time to take care of their children, including feeding them properly and adequately which therefore could lead to meeting their MAD. In addition, if there is access to drinking water source this could also encourage households to use water for on-plot gardening, which can improve dietary diversity. Furthermore, if there is access to drinking water, time spent to fetching water could be invested in additional productive activities that can augment economic and educational empowerment of mothers.

In this study exposure to media was also a significant predictor of feeding minimum acceptable diet to the child. Families who have exposure to media had increased odds of feeding minimum acceptable diet to their child than those who have no media exposure. This is supported by studies conducted in south Asia and India which reported low feeding practice of minimum acceptable diet for the children in the families who have no media exposure [24, 26]. This might be pointing to the influence of media on behavioral change to improve the minimum acceptable diet through enhancing mother’s knowledge on feeding a minimum acceptable diet to their children.

From the community level factors geographic region of the study participants was found to be statistically significant predictor of feeding minimum acceptable diet in Ethiopia. Children from city dwellers and agrarian dominant had more odds to be fed minimum acceptable diet than children from pastoralist dominant region. This might be due to Ethiopia is large country with diverse cultures, religions reflected by different food habits and traditional practices [21]. This could also be linked with less food production and higher levels of poverty.

Furthermore, the current study found that; even though most variation on feeding the child minimum acceptable diet was explained by individual-level factors, some of the variation was also explained by unmeasured community-level factors. The random effects of the community-level model were significant in explaining feeding the child minimum acceptable diet, even it reduced, from 39.7% in null model to 27% in the full model. This indicates feeding the child minimum acceptable diet was explained by both individual and community-level factors. The efforts of improving the practice of MAD in Ethiopian should address both factors operating at individual and community level.

## Conclusion

The proportion of young children aged between 6–23 months receiving minimum acceptable diet was very low (6.1%). This study also showed that this low proportion of minimum acceptable diet among 6-23 months age children is determined by number of individual and community level factors such as age of the child, mother’s occupation, father’s education level, and media exposure, access of household drinking water and region of residence. Therefore, interventions to improve MAD practice should not only be implemented at the individual level but also be tailored to the community context. Utilizing media to promote feeding young children with minimum acceptable diet through enhancing mother’s knowledge on child feeding practices and advocacy for appropriate complementary feeding, particularly on meeting MAD and nutrition education and social and behavior change interventions should be strengthened targeting the infants and children aged 6–8 months and unemployed mothers.

## Acknowledgments

Our sincere and deepest gratitude goes to Mekelle University, College of Health Sciences and MU-NMBU for financial support of this study. We are also very grateful to Measure DHS for making the data freely available and we would like to extend our heartfelt thanks and appreciation to family members of Mrs.Aberash Abay (Haset Beyene, Nobel Beyene, Lidiya Beyene and Beyene Meressa) and friends for their endless support and providing the required courage during the research work.

## Author Contributions

**Conceptualization:** Aberash Abay, Dejen Yemane, Abate Bekele, Beyene Meressa

**Formal analysis:** Aberash Abay, Dejen Yemane, Abate Bekele, Beyene Meressa

**Funding acquisition:** Aberash Abay

**Investigation:** Aberash Abay, Dejen Yemane, Abate Bekele

**Methodology:** Aberash Abay, Dejen Yemane, Abate Bekele, Beyene Meressa

**Project administration:** Aberash Abay, Dejen Yemane, Abate Bekele

**Supervision:** Dejen Yemane, Abate Bekele, Beyene Meressa

**Validation:** Aberash Abay, Dejen Yemane, Abate Bekele, Beyene Meressa.

**Writing original draft:** Aberash Abay, Dejen Yemane, Abate Bekele, Beyene Meressa.

**Writing review & editing:** Aberash Abay, Dejen Yemane, Abate Bekele, Beyene Meressa

